# h-channels contribute to divergent electrophysiological properties of supragranular pyramidal neurons in human versus mouse cerebral cortex

**DOI:** 10.1101/312298

**Authors:** Brian E Kalmbach, Anatoly Buchin, Jeremy A Miller, Trygve E Bakken, Rebecca D Hodge, Peter Chong, Rebecca de Frates, Kael Dai, Ryder P. Gwinn, Charles Cobbs, Andrew L Ko, Jeffrey G Ojemann, Daniel L Silbergeld, Christof Koch, Costas A. Anastassiou, Ed Lein, Jonathan T Ting

## Abstract

Gene expression studies suggest that differential ion channel expression contributes to differences in rodent versus human neuronal physiology. We tested whether h-channels more prominently contribute to the physiological properties of human compared to mouse supragranular pyramidal neurons. Single cell/nucleus RNA sequencing revealed ubiquitous HCN1-subunit expression in excitatory neurons in human, but not mouse supragranular layers. Using patch-clamp recordings, we found stronger h-channel-related membrane properties in supragranular pyramidal neurons in human temporal cortex, compared to mouse supragranular pyramidal neurons in temporal association area. The magnitude of these differences depended upon cortical depth and was largest in pyramidal neurons in deep L3. Additionally, pharmacologically blocking h-channels produced a larger change in membrane properties in human compared to mouse neurons. Finally, using biophysical modeling, we provided evidence that h-channels promote the transfer of theta frequencies from dendrite-to-soma in human L3 pyramidal neurons. Thus, h-channels contribute to between-species differences in a fundamental neuronal property.

## Introduction

The development and gross anatomical organization of the mammalian cerebral cortex is stereotyped across species (Angevine and Sidman, 1961; DeFelipe, 2011; Rakic, 1974). However, while all mammals possess a hexalaminar cortex comprising diverse neuronal cell types, the primate cortex has undergone dramatic evolutionary expansion, especially in layers (L) 2 and 3 (DeFelipe, 2011). Consequently, in contrast to rodent, L2 and 3 in human neocortex are easily distinguishable from each other and further sublaminar distinctions within layer 3 can be made based upon the size and organization of pyramidal neurons (DeFelipe, 2011; von Economo and Koskinas, 2007; Hill and Walsh, 2005; Molnár et al., 2014; Rakic, 2009). This evolutionary expansion and stratification suggests that there is functional specialization of neuronal properties within the supragranular layers of human neocortex.

Cross-species comparisons of supragranular pyramidal neuron properties indeed suggest that such neuronal specializations exist. For example, supragranular pyramidal neurons are larger and display more complex dendritic branching in human compared to the mouse (Deitcher et al., 2017; Mohan et al., 2015). Additionally, there appear to be differences in the passive membrane properties of human neurons that may compensate for the filtering of electrical signals that would occur along such extensive dendritic arbors (Eyal et al., 2016). These differences may contribute to the unique cable properties of human pyramidal neuron dendrites and the enhanced ability of human neurons to track high frequency synaptic input (Eyal et al., 2014; Testa-Silva et al., 2014).

While these differences in passive neuronal properties are notable, surprisingly little is known about how differential ion channel expression contributes to differences in active membrane properties between human versus rodent neurons. Voltage-gated ion channels shape a neuron’s subthreshold integrative properties and endow it with the ability to generate non-linear, regenerative events, including axonal action potentials and dendritic spikes (Bean, 2007; Johnston and Narayanan, 2008; Koch, 2004; Reyes, 2001; Stuart and Spruston, 2015). In this way, ion channels are prime contributors to specialized neuronal function. Intriguingly, large-scale cross-species comparisons of gene expression have revealed differences in the laminar expression of several ion channel-associated genes between mouse and human cortex (Zeng et al., 2012). Specifically, RNA for HCN1, a major pore-forming subunit of h-channels (Robinson and Siegelbaum, 2003) is differentially expressed in human versus mouse neocortex (Zeng et al., 2012). The hyperpolarization-activated non-specific cation current, I_h_, which is carried by h-channels, greatly shapes a neuron’s subthreshold integrative properties (Magee, 1998; Robinson and Siegelbaum, 2003; Williams and Stuart, 2000).

Here, we use single nucleus transcriptomics and *in vitro* slice physiology to provide evidence that h-channels contribute to the membrane properties of supragranular pyramidal neurons in human cortex more so than in mouse cortex. We then use a biophysical model to provide insight into how the presence of h-channels affects the integrative properties of human supragranular pyramidal neurons. Our findings implicate a species-specific role for h-channels in dendritic integration of synaptic input that is most pronounced in deep L3 pyramidal neurons of human temporal cortex, which may generally represent an important evolutionary adaptation for very large pyramidal neurons in the human neocortex.

## Experimental Procedures

### Human surgical specimens

Surgical specimens were obtained from local hospitals (Harborview Medical Center, Swedish Medical Center and University of Washington Medical Center) in collaboration with local neurosurgeons. All patients provided informed consent and experimental procedures were approved by hospital institute review boards before commencing the study. The bulk of data included in this study were obtained from tissue from 10 patients with temporal lobe epilepsy with a mean age of 38.10 ± 15.67. Four patients were male and 6 were female. Additionally, data were obtained from tissue from one patient who had undergone deep tumor resection from the temporal lobe. Tissue obtained from surgery was distal to the core pathological tissue and was deemed not to be of diagnostic value. Specimens were placed in a sterile container filled with an artificial cerebral spinal fluid (aCSF) composed of (in mM): 92 with N-methyl-D-glucamine (NMDG), 2.5 KCl, 1.25 NaH_2_PO_4_, 30 NaHCO_3_, 20 4-(2-hydroxyethyl)-1-piperazineethanesulfonic acid (HEPES), 25 glucose, 2 thiourea, 5 Na-ascorbate, 3 Na-pyruvate, 0.5 CaCl_2_·4H_2_O and 10 MgSO_4_·7H_2_O. The pH of the NMDG aCSF was titrated to pH 7.3–7.4 with concentrated hydrochloric acid and the osmolality was 300-305 mOsmoles/Kg. The solution was pre-chilled to 2-4°C and thoroughly bubbled with carbogen (95% O_2_/5% CO_2_) prior to collection.

Surgical specimens were quickly transported from the surgical site to the laboratory while continuously bubbled with carbogen (transportation time: 10-40 minutes).

### Acute brain slice preparation

To ensure that the dendrites of pyramidal neurons were relatively intact, human surgical specimens were trimmed and mounted such that the angle of slicing was perpendicular to the pial surface. 350 μm thick slices were sectioned on a Compresstome VF-200 (Precisionary Instruments) using the NMDG protective recovery method (Ting et al., 2014) and either a zirconium ceramic blade (EF-INZ10, Cadence) or a sapphire knife (93060, Electron Microscopy Sciences). The slicing solution was the same as used for transport from the hospital to the laboratory. After all sections were obtained, slices were transferred to a warmed (32-34° C) initial recovery chamber filled with NMDG aCSF under constant carbogenation. After 12 minutes, slices were transferred to a Brain Slice Keeper-4 holding chamber (Automate Scientific) containing an aCSF solution made of (in mM): 92 NaCl, 2.5 KCl, 1.25 NaH_2_PO_4_, 30 NaHCO_3_, 20 HEPES, 25 glucose, 2 thiourea, 5 Na-ascorbate, 3 Na-pyruvate, 2 CaCl_2_·4H_2_O and 2 MgSO_4_·7H_2_O continuously bubbled with 95/5 O_2_/CO_2_. Slices were held in this chamber at room temperature for 1-48 hours before transfer to the recording chamber for patch clamp recording.

All procedures involving mice were approved by the Institutional Animal Care and Use Committee. Mouse brain slices were prepared in largely the same fashion as human slices. Four male and 6 female mice, 44-61 days old (49.8 ± 5.59), were deeply anesthetized by intraperitoneal administration of Advertin (20 mg/kg IP) and were perfused through the heart with NMDG aCSF (bubbled with carbogen). Coronal slices containing the temporal association area (TeA) were prepared as described for human with the exception that slices were 300 μm rather than 350 μm thick.

### Patch clamp recordings

Slices were placed in a submerged, heated (32-34° C) recording chamber that was continually perfused (3-4 mL/min) with aCSF under constant carbogenation and containing the following (in mM): 119 NaCl, 2.5 KCl, 1.25 NaH_2_PO_4_, 24 NaHCO_3_, 12.5 glucose, 2 CaCl_2_4H_2_O and 2 MgSO_4_·7H_2_O (pH 7.3-7.4). Slices were viewed with an Olympus BX51WI microscope and infrared differential interference contrast optics and a 40x water immersion objective. Patch pipettes (3-6 MΩ) were pulled from borosilicate glass using a horizontal pipette puller (P1000, Sutter Instruments). The pipette solution for all experiments contained the following (in mM): 130 K-gluconate, 10 HEPES, 0.3 EGTA, 4 Mg-ATP, 0.3 Na_2_-GTP and 2 MgCl_2_. The pipette solution also contained 5% biocytin and 20 μM Alexa 594. These were included to ensure that the apical dendrite reached the pial surface. Alexa filled cells were visualized only upon termination of the recording using a 540/605 nm excitation/emission filter set. The theoretical liquid junction potential was calculated to be -13 mV and was not corrected

Whole cell somatic recordings were acquired using a Multiclamp 700B amplifier and PClamp 10 data acquisition software (Molecular Devices). Electrical signals were digitized (Axon Digidata 1550B) at 20-50 kHz and filtered at 2-10 kHz. Upon attaining whole-cell current clamp mode, the pipette capacitance was compensated and the bridge was balanced. Access resistance was monitored throughout the recording and was 8-25 MΩ. Recordings were terminated if access resistance exceeded 25 MΩ. ZD7288 (10 μM; Tocris) was prepared from frozen concentrated stock solutions and diluted in recording aCSF.

### Data analysis and statistical testing

Data were analyzed using custom analysis scripts written in Igor Pro (Wavemetrics). All measurements were made at resting membrane potential and, in a subset of experiments, at a common potential of -65 mV. Input resistance (R_n_) was calculated from the linear portion of the current-voltage relationship generated in response to a series of 1s current injections (–150 to +50 pA, 20 or 50 pA steps). The maximum and steady state voltage deflections were used to determine the maximum and steady state of R_N_, respectively. Voltage sag was defined as the ratio of maximum to steady-state R_N_. Rebound slope was calculated from the slope of the rebound amplitude as a function of steady-state membrane potential. Resonance was determined from the voltage response to a constant amplitude sinusoidal current injection that linearly increased in frequency from 1-14 or 15 Hz over 15 s. The impedance amplitude profile (ZAP) was constructed from the ratio of the fast Fourier transform of the voltage response to the fast Fourier transform of the current injection. The frequency corresponding to the peak impedance (Z_ma_χ) was defined as the resonant frequency. The 3dB cutoff was calculated as the frequency at which the ZAP profile attenuated to a value of (√1/2) Z_max_. Action potentials (APs) were elicited in response to increasing amplitude, 1s direct current injections (50-750 pA, 50 pA steps).

Statistical analyses and plotting were performed using Prism (Graphpad). Data are presented in the text as mean ± SEM or R^2^ values. Between-subject ANOVA, mixed factors ANOVA, two-sample Kolmogorov-Smirnov test and post hoc t-tests were used to test for statistical differences between groups.

Bonferroni correction was used to correct for multiple comparisons. Pearson’s product moment correlation was used to test for statistically significant correlations between variables.

### Biophysical model

The morphological reconstruction (Figure 7) was generated using methods previously described (Allen Institute for Brain Science, 2016; Gouwens et al., 2018). Briefly, the tissue slice was stained via diaminobenzidine reaction to elucidate biocytin filled cells. Z-stacks of the cell were imaged at 63x magnification on a Zeiss Axio Imager 2. Reconstruction was manually performed using Vaa3D software to create accurate full neuron representations of the soma, dendrites, and axon saved in the SWC format (Peng et al., 2010).

The single neuron model was simulated using the Neuron 7.4 simulator (https://www.neuron.yale.edu/neuron/) in combination with the Brain Modeling Toolkit (https://github.com/AllenInstitute/bmtk). The morphology of the simulated neuron was adopted from the SWC file generated in the morphology reconstruction process. The dendritic tree was discretized using 20 μm spatial steps and corresponding cylindrical compartments were generated for all compartments. The model was simulated using temporal steps of 0.1 ms. We insured that further reduction of spatial and temporal discretization steps did not provide quantitatively different results. Passive membrane parameters were adjusted for axonal, somatic, apical and basal dendrites compartments. Leak conductance, capacitance, leak reversal potential and axial resistance were fitted to the subthreshold responses of a human pyramidal neuron in response to the family of somatic current injections used to estimate passive parameters using genetic algorithms from DEAP library (https://github.com/DEAP/deap). The best parameters to fit the subthreshold responses were: 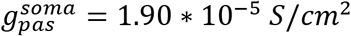; 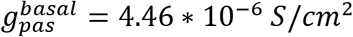; 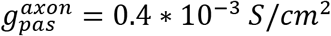; 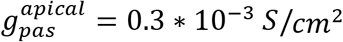; 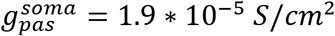; 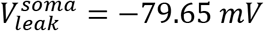; 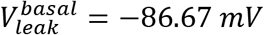; 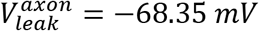; 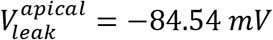; 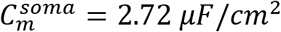; 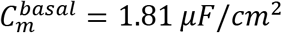; 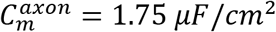; 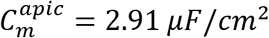; 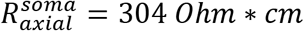; 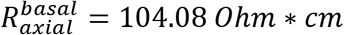; 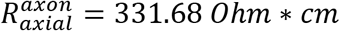; 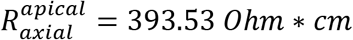.

The kinetic parameters of I_h_ were adopted from (Kole et al., 2006) and m_alp_ha function was shifted by -20 mV to match the junction potential estimation from the experimental data. I_h_ and the corresponding HCN conductance was uniformly distributed in somatic, axonal and dendritic compartments, *g_HCN_* = 0.01 * 10^−3^ *S*/*cm*^2^. The maximal conductance (*g_HCN_*) for I_h_ was adjusted such that the difference in R_N_ between the I_h_(+) and I_h_(-) models matched the difference in R_N_ observed in experiments after the blockade of I_h_ current by ZD7288. The model possessed no other active conductances. The model code is available on github page https://github.com/AllenInstitute/***.

Synaptic conductances (10 nS) were simulated using AMPA-like kinetics: T_rise_ = 1 5 ms, T_decay_ = 3 ms, reversal potential = 0 mV (Andrasfalvy and Magee, 2001; Jonas and Sakmann, 1992; Spruston et al., 1995). For simulations involving random activation of 1000 synaptic inputs, the synaptic conductance of each synapse was scaled to 0.001 mS/cm^2^. Three random seeds were instantiated using Python Numpy to generate temporal permutations of synaptic population activation (total simulation duration: 30 s). All results of simulations were stored in NWB file format. Synapses were activated by a Poisson process with 4 Hz rate generated using Numpy expovariate() function. All simulations were performed on a MacBook Pro Retina with 2.8 GHz Intel core i7processor with 16 GB DDR3 RAM running Mac OS 10.13.3 Sierra.

## Results

To gain initial insight into whether human supragranular pyramidal neurons express h-channels, we utilized the existing online Allen Cell Types Database (www.celltypes.brain-map.org; methods are included on website) consisting of single-nucleus RNA sequencing (n=15,928 nuclei) from human postmortem brain specimens. Briefly, this method involved layer dissections of thin Nissl-stained human temporal cortex section, neuronal nuclei staining (NeuN) and Fluorescence-activated cells sorting (FACS) isolation, followed by Smart-seq v4 based library preparation and single-cell deep (2.5 million reads/cell) RNA-Seq. *GAD1* and *SLC17A7* expression were used to delineate inhibitory and excitatory neurons, respectively. For a comparison, in mouse we analyzed a previously published single cell RNA sequencing dataset obtained from mouse primary visual cortex (Tasic et al., 2016). We examined the expression levels of the four pore forming subunits (*HCN1-4*) of h-channels as well as *PEX5L* which codes for a protein (Trip8b) involved in modulating h-channel function and dendritic enrichment (Lewis et al., 2009; Robinson and Siegelbaum, 2003; Santoro, 2004).

In human temporal cortex, *HCN1* and *PEX5L* were prominently expressed in excitatory neurons in the supragranular and infragranular layers, whereas *HCN2-4* subunits were much less abundant (Figure 1A). The average expression of *HCN1* was approximately equal (1.1 fold higher) in supragranular compared with infragranular excitatory neurons. In contrast, in the mouse neocortex *HCN1* and *PEX5L* were abundant in infragranular, but not supragranular excitatory neurons (Figure 1B). The average expression of *HCN1* was 3 fold higher in infragranular compared with supragranular excitatory neurons. Furthermore, *HCN2-4* expression was relatively very low in excitatory neurons across the layers. Consistent with these observations, *HCN1* expression was observed in the supragranular and infragranular layers of human temporal cortex by ISH (Figure 1C; data obtained from the Allen Brain Atlas, human.brain-map.org), whereas in mouse it was seen most conspicuously in L5 in multiple brain regions, including primary visual cortex and temporal association area (TeA; Figure 1D; data obtained from Allen Brain Atlas, mouse.brain-map.org). Notably, *HCN1* expression was relatively high in interneuron populations in mouse and human cortical tissue (Figures 1A and B), suggesting that the scattered labeling in the supragranular layers of mouse cortex via ISH represent interneurons (Figure 1D). Together these data suggest that there is widespread expression of h-channels, especially those containing HCN1 subunits, in human, but not mouse, supragranular pyramidal neurons.

**Figure 1.**
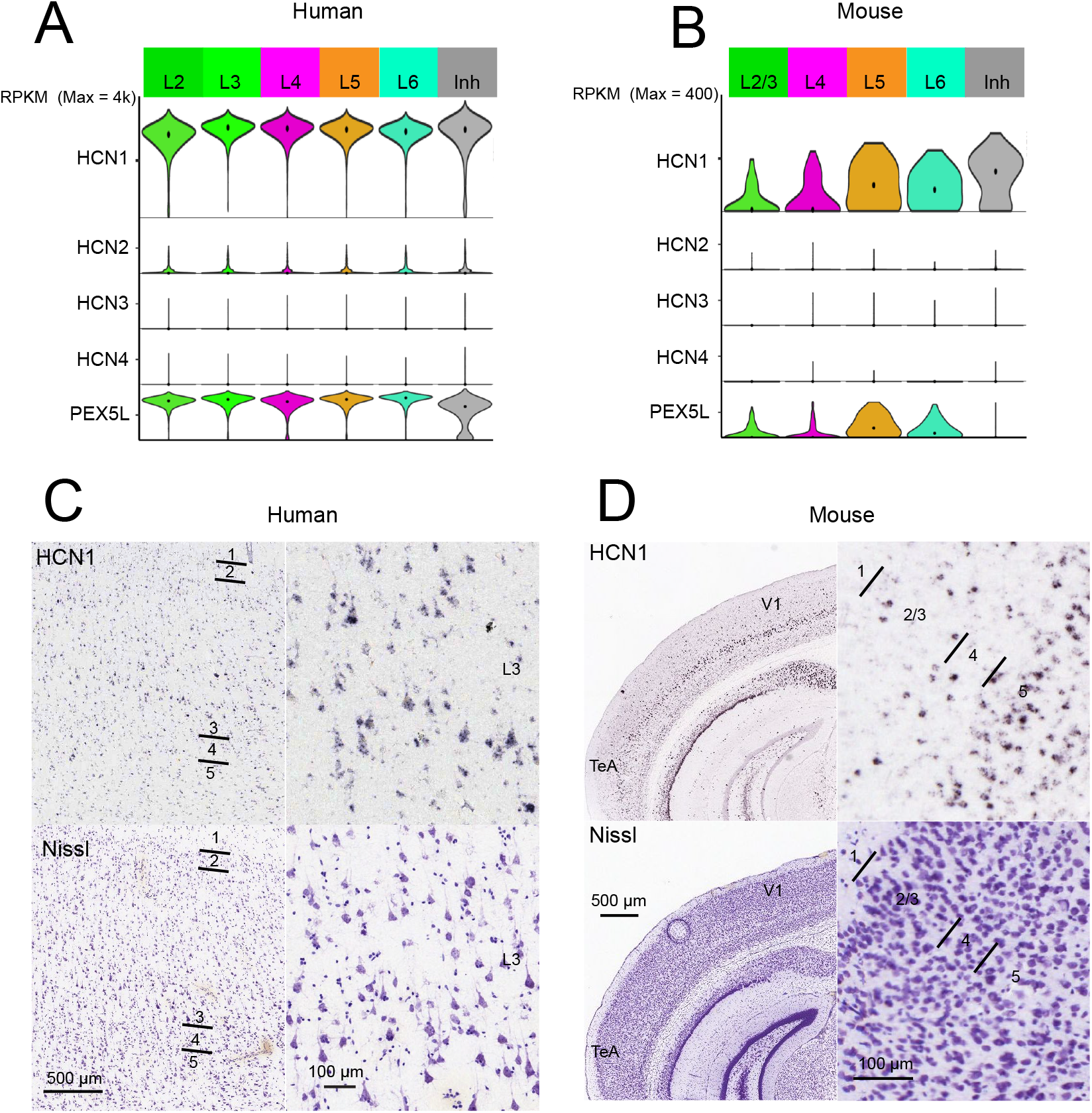
– *HCN1-4* RNA expression in human versus mouse neocortex. **A)** Single nucleus HCN channel subunit mRNA expression in excitatory neurons from human temporal cortex arranged by layer. Violin plots represent distribution of mRNA expression on a log scale with a maximum reads per kilobase per million mapped reads (RPKM) value of 4000. For reference, single nucleus HCN channel subunit mRNA expression in inhibitory neurons across all layers is also shown. There are approximately equal levels of *HCN1* expression in excitatory neurons across the cortical layers. **B)** Single cell HCN channel subunit mRNA expression in excitatory neurons from mouse visual cortex arranged by layer. Violin plots represent distribution of mRNA expression on a log scale with a maximum RPKM value of 400, ten times lower than for human cells. Lower levels of h-channel-related gene expression were observed in excitatory neurons in the supragranular compared with infragranular layers of mouse primary visual cortex. Robust *HCN1* expression was also observed in inhibitory neurons. **C)** ISH of *HCN1* in human temporal cortex at low magnification (left column, with near adjacent Nissl stained section for layer identification) and high magnification in deep L3. **D)** ISH of *HCN1* in mouse neocortex at low magnification (left column, with near adjacent Nissl stained section) and high magnification of TeA.

### Physiological evidence for differences in I_h_

To allow for direct between-species comparisons of pyramidal neuron membrane properties as a function of somatic distance from the pial surface, we first examined differences in the gross cytoarchitecture of mouse and human cortex. For these purposes we chose the TeA of mouse cortex because it has been used in previous studies as a comparator for the middle temporal cortex typically resected from human patients (Eyal et al., 2016; Mohan et al., 2015; Testa-Silva, 2010; Wang et al., 2015). We performed DAPI staining on thick (350 μm) sections of mouse temporal association (4 slices from 4 mice) and human temporal (6 slices from 6 patients) cortex (Figure S1A). In both species we observed a sharp increase in cell density marking the boundary between L1 and L2 (mouse 151 ± 14 μm, human 276 ± 12 μm from pial surface). Below L2 was a sparser region of cells (L3) followed by a tight band of densely packed cells (L3/L4 boundary, mouse 470 ± 6 μm, human 1469 ± 34 μm from pial surface). Notably, the distance from the pial surface to the L3/L4 border ranged from 1292-1637 μm in human sections and 453-508 μm in mouse sections. L4 was followed by a decrease in cell density, marking L5 (mouse 599 ± 10 μm from pial surface, human 1736 ± 37 μm from pial surface). Finally, the bottom of L6 was 2963 ± 97 μm and 1252 ± 29 μm from the pial surface in human and mouse neocortex, respectively. Thus, L2/3 represents ~40% and ~25% of the total thickness of human temporal gyrus and mouse TeA, respectively. These observations are consistent with previous reports that illustrate the large expansion of supragranular cortex in the human cortical column relative to mouse (DeFelipe, 2011; von Economo and Koskinas, 2007).

We performed whole cell patch clamp recordings from supragranular pyramidal neurons with cell bodies located throughout the entire depth of L2 and L3 (sample biocytin fills in Figures S1B and C; human n=55 cells between 350-1600 μm, mouse n=39 cells between 162-465 μm from the pial surface). Using IR-DIC optics, L3 was easily distinguishable from L4 both in terms of cell size and density; deep L3 consisted of sparse, large pyramidal neurons whereas L4 consisted of more densely packed granular-appearing cells. Morphological differences between mouse and human supragranular pyramidal neurons have been extensively detailed elsewhere, thus we did not pursue them further (Deitcher et al., 2017; Mohan et al., 2015).

We compared I_h_-related membrane properties in human versus mouse pyramidal neurons. h-channels are open at relatively hyperpolarized potentials and thus contribute to a neuron’s input resistance (R_N_) and resting membrane potential (RMP; Magee, 1998; Robinson and Siegelbaum, 2003). As an initial test for differences in the functional expression of h-channels, we measured the RMP and R_N_ of pyramidal neurons throughout the depth of mouse and human supragranular cortex. Sample voltage responses obtained from a superficial and deep supragranular pyramidal neuron from both species are shown in Figures 2A and B. In both mouse and human temporal cortex, we observed a positive correlation between RMP and somatic distance from the pial surface, such that the most depolarized neurons were found deep in L3 (mouse r^2^ = 0.35, p<0.001; human r^2^ = 0.19, p<0.001; Figures 2C-E). In contrast, the R_N_ of mouse neurons increased as a function of somatic distance from pial surface (0.34, p<0.001; Figures 2C and E) whereas the R_N_ of human neurons decreased (r^2^= 0.31, p<0.001; Figures 2D and E). Thus, in human temporal cortex, the neurons with the lowest input resistance were found deep in L3 whereas in mouse cortex they were found superficially, near the L1/2 border (Figure 2E). For human temporal cortex, these general observations were replicated in a subset of experiments in which R_N_ was measured at a common membrane potential of -65 mV (Figure S2; n = 43, r^2=^0.32, p < 0.001). In contrast, in mouse there was no correlation between R_N_ and somatic distance from pia in a subset of experiments performed at -65 mV (Figure S2; n = 24, r^2=^0.02, p= 0.53). Notably, the observation that R_N_ increases as a function of somatic depth from the pial surface in mouse temporal association cortex is consistent with a previous report from mouse prefrontal cortex (Routh et al., 2017). Thus, the depth-dependence of R_N_ observed here might be a hallmark of mouse L2/3 regardless of cortical region.

**Figure 2.**
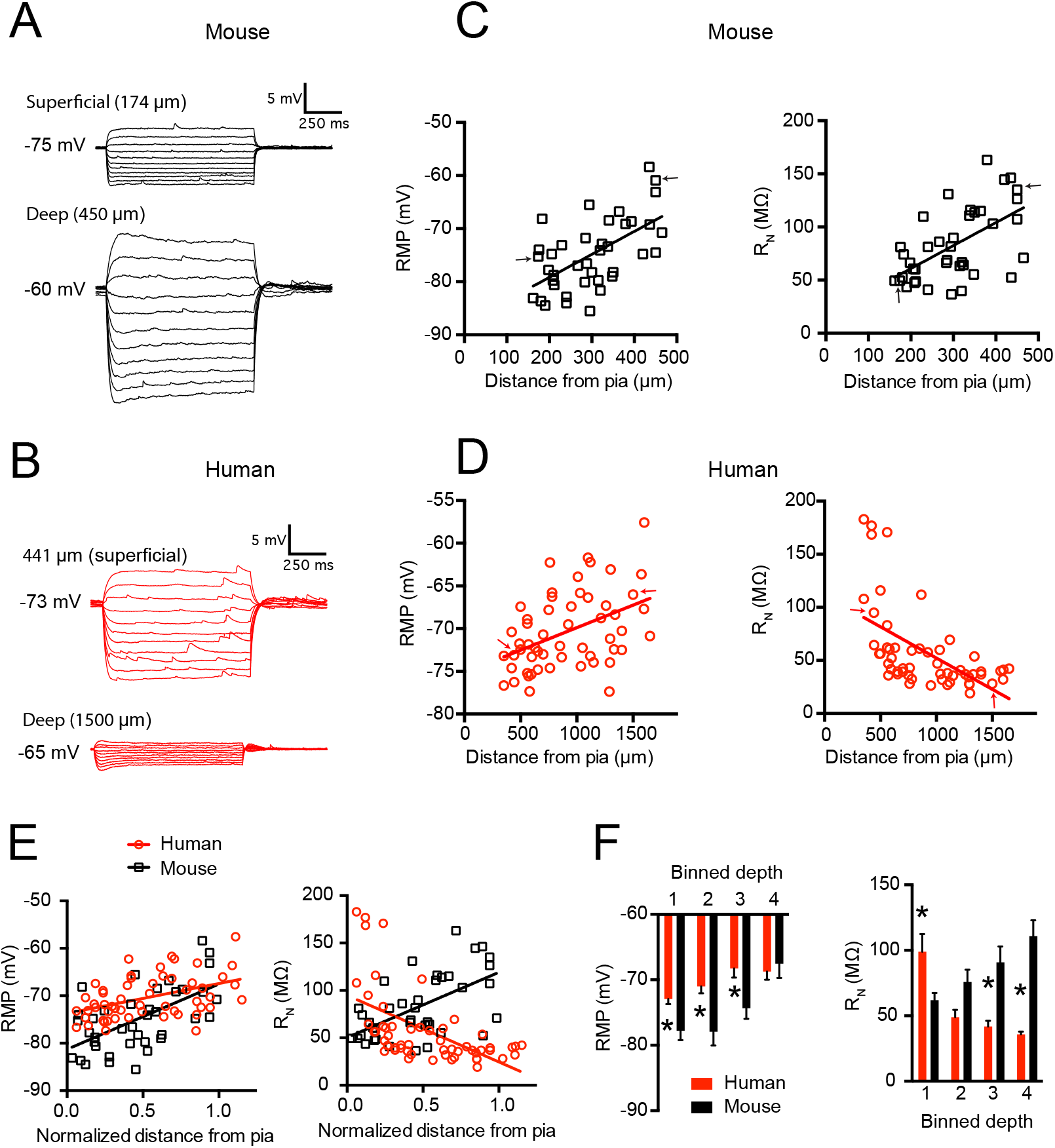
– *HCN1-4* RNA expression in human versus mouse neocortex. Human and mouse supragranular pyramidal neurons display different subthreshold membrane properties. Example voltage sweeps obtained from a superficial and deep supragranular pyramidal neuron in response to a family of hyperpolarizing and depolarizing current injections in **A)** mouse temporal association area and **B)** human middle temporal gyrus. **C)** In mouse cortex, resting membrane potential and input resistance increase as a function of somatic distance from pia. **D)** In the supragranular layers of human middle temporal gyrus resting potential increases, but input resistance decreases as function of somatic distance from pia. Arrows correspond to sample voltage sweeps in A & B **E)** Resting potential and input resistance in mouse versus human cortex as a function of normalized somatic position in supragranular cortex. **F)** Data were also binned into four groups based on the normalized distance of the soma from pia, where 1 is the most superficial quadrant and 4 is the deepest. * p < 0.0125, mouse versus human post-hoc t-test with Bonferroni correction.

To make direct comparisons between mouse and human pyramidal neuron properties, we binned the data into quarters based on normalized soma distance from the pial surface to the border of L3 and L4. Human neurons were more depolarized than mouse neurons throughout the first three quarters of supragranular cortex (p = 0.03; ANOVA followed by post-hoc comparisons;

Figure 2F). Furthermore, human neurons displayed a higher R_N_ in the most superficial portion of supragranular cortex and lower R_N_ in the lower half of supragranular cortex (p < 0.001; ANOVA followed by post-hoc comparisons; Figure 2F). These data demonstrate significant distance-dependent differences in the properties of human versus mouse supragranular pyramidal neurons.

In addition to contributing to R_N_, I_h_ contributes to several unique membrane properties. Specifically, I_h_ is associated with a characteristic voltage “sag” upon membrane hyperpolarization and a rebound potential upon release from hyperpolarization (Figures 3A and B). These voltage responses reflect the activation and deactivation kinetics of h-channels (Robinson and Siegelbaum, 2003). While in the supragranular layers of mouse temporal association area these I_h_-related membrane properties were largely absent, we did observe a few neurons deep in L3 that displayed modest amounts of sag and rebound (sag r^2^ = 0.14, p = 0.02, rebound sag r^2^ = 0.26, p < 0.001; Figure 3C). In contrast, sag and rebound were apparent in many human pyramidal neurons throughout the supragranular layers and were positively correlated with somatic depth from the pial surface (sag r^2^ = 0.18, p = 0.001, rebound r^2^ = 0.20, p < 0.001; Figure 3D). These response properties are plotted as a function of normalized depth from pia to the border of L3 and L4 and illustrate the marked differences in I_h_-related properties between mouse and human supragranular pyramidal neurons (Figure 3E). These general observations were replicated in a subset of experiments performed at -65 mV (Figures S2 and S3; mouse n=24, sag r^2^ = 0.01, p = 0.65, rebound r^2^ = 0.35, p = 0.002; human n=43, sag r^2^ = 0.14, p = 0.01, rebound r^2^ = 0.22, p < 0.001). Finally, direct between-species comparisons revealed that human neurons possessed more sag and rebound at all levels of supragranular cortex compared to their mouse counterparts (Figure 3F; p < 0.001; ANOVA followed by post-hoc comparisons).

**Figure 3.**
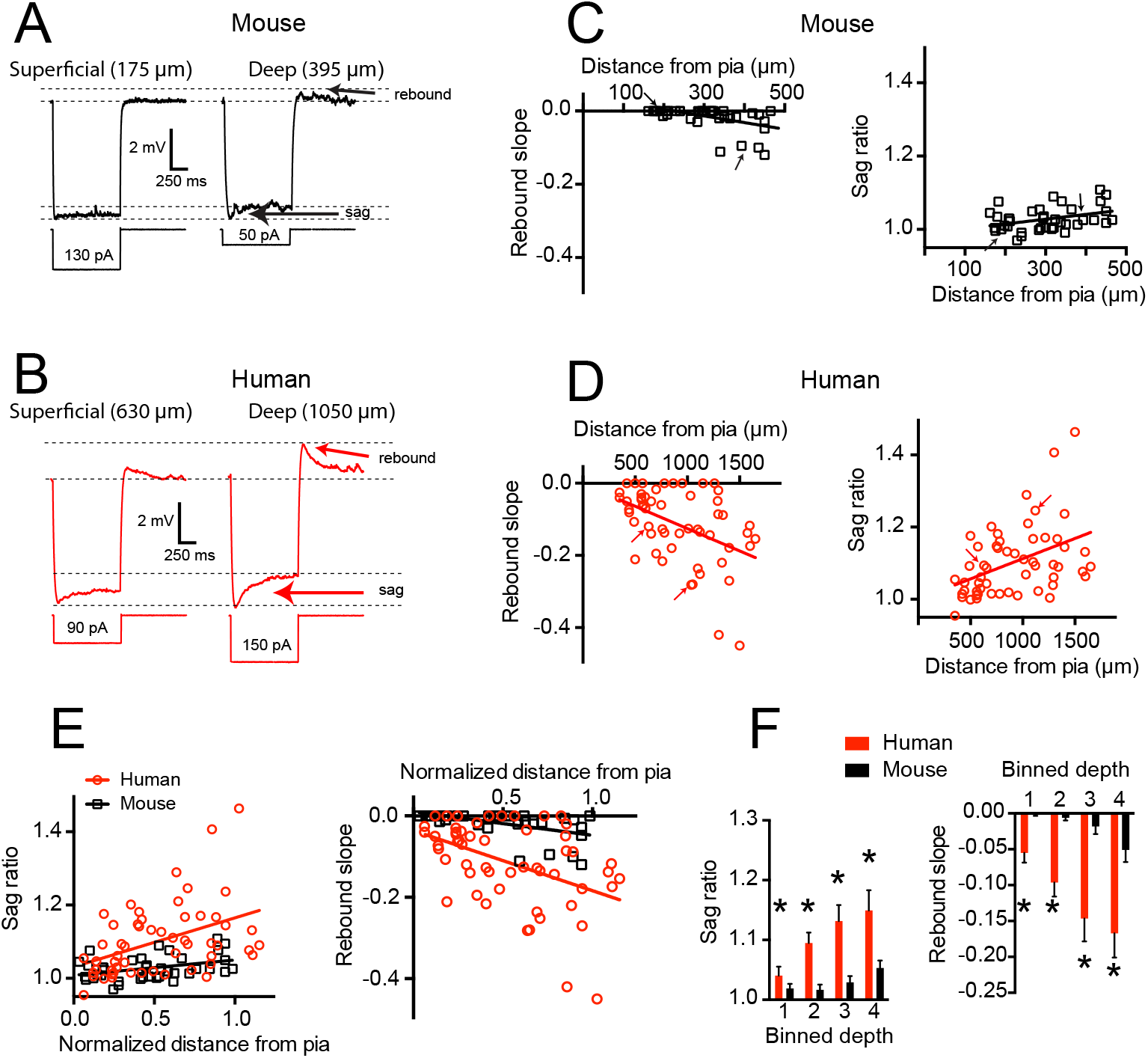
– I_h_-related membrane properties are more pronounced in human compared with mouse supragranular pyramidal neurons. **A)** Example voltage sweeps were obtained from current injections that yielded ~ 6 mV hyperpolarization in **A)** mouse and **B)** human supragranular pyramidal neurons. Arrows denote voltage sag and rebound potentials associated with the presence of I_h_. **C)** Mouse neurons display little voltage sag or rebound from hyperpolarization in response to hyperpolarizing current injections. **D)** In contrast, rebound and sag were prominent in human supragranular cortex, especially in deep layer 3. Arrows correspond to sample voltage sweeps in A & B. **E)** Sag and rebound in mouse and human cortex as a function of normalized somatic position in supragranular cortex. **F)** As before, data were also binned into four groups. * p < 0.001 mixed factor ANOVA effect of species.

The slow activation and deactivation kinetics of h-channels contribute greatly to the filtering properties of a neuron. Specifically, I_h_ contributes to membrane resonance in the ~2-7 Hz range (Dembrow et al., 2010; Hutcheon et al., 1996; Kalmbach et al., 2013; 2017; 2015; Narayanan and Johnston, 2007; Nolan et al., 2004; Ulrich, 2002). To test for differences in the subthreshold filtering properties of human versus mouse pyramidal neurons, we measured the response of pyramidal neurons throughout the depth of supragranular cortex to a chirp stimulus. In addition to measuring the resonant frequency of neurons, we also calculated the 3 dB cutoff as a way to quantify differences in low-pass filtering. Sample voltage responses, ZAPs and normalized frequency response curves for superficial and deep mouse and human neurons are shown in Figures 4A and B. Mouse pyramidal neurons were largely non-resonant at either RMP or -65 mV, although there were a few neurons located deep in L3 that showed modest resonance (Figures 4C and S3; r^2^ = 0.13, p = 0.02; -65 mV r^2^ = 0.01, p = 0.59). Additionally, the 3dB cutoff of mouse neurons was negatively correlated with somatic depth from pia at RMP (Figure 4C, r^2^ = 0.12, p = 0.03), but not at - 65 mV (Figure S3; r^2^ = 0.01, p = 0.99) indicating that the filtering properties of deep supragranular pyramidal neurons were more low-pass than superficial ones at RMP. In contrast, many human pyramidal neurons displayed membrane resonance in the ~2-5 Hz range. Indeed, resonance was positively correlated with somatic depth from pia in human supragranular cortex when measured at RMP (Figure 4D; r^2^ = 0.13, p = 0.007) and -65 mV (Figure S2 1; r^2^ = 0.32, p < 0.001). In contrast to mouse pyramidal neurons, the 3dB cutoff of human pyramidal neurons was positively correlated with somatic depth from pia (Figure 4D; RMP, r^2^ = 0.12, p = 0.01; Figure S2; -65 mV r^2^ = 0.22, p = 0.001), such that the filtering properties of superficial neurons were distinct from deep neurons.

**Figure 4.**
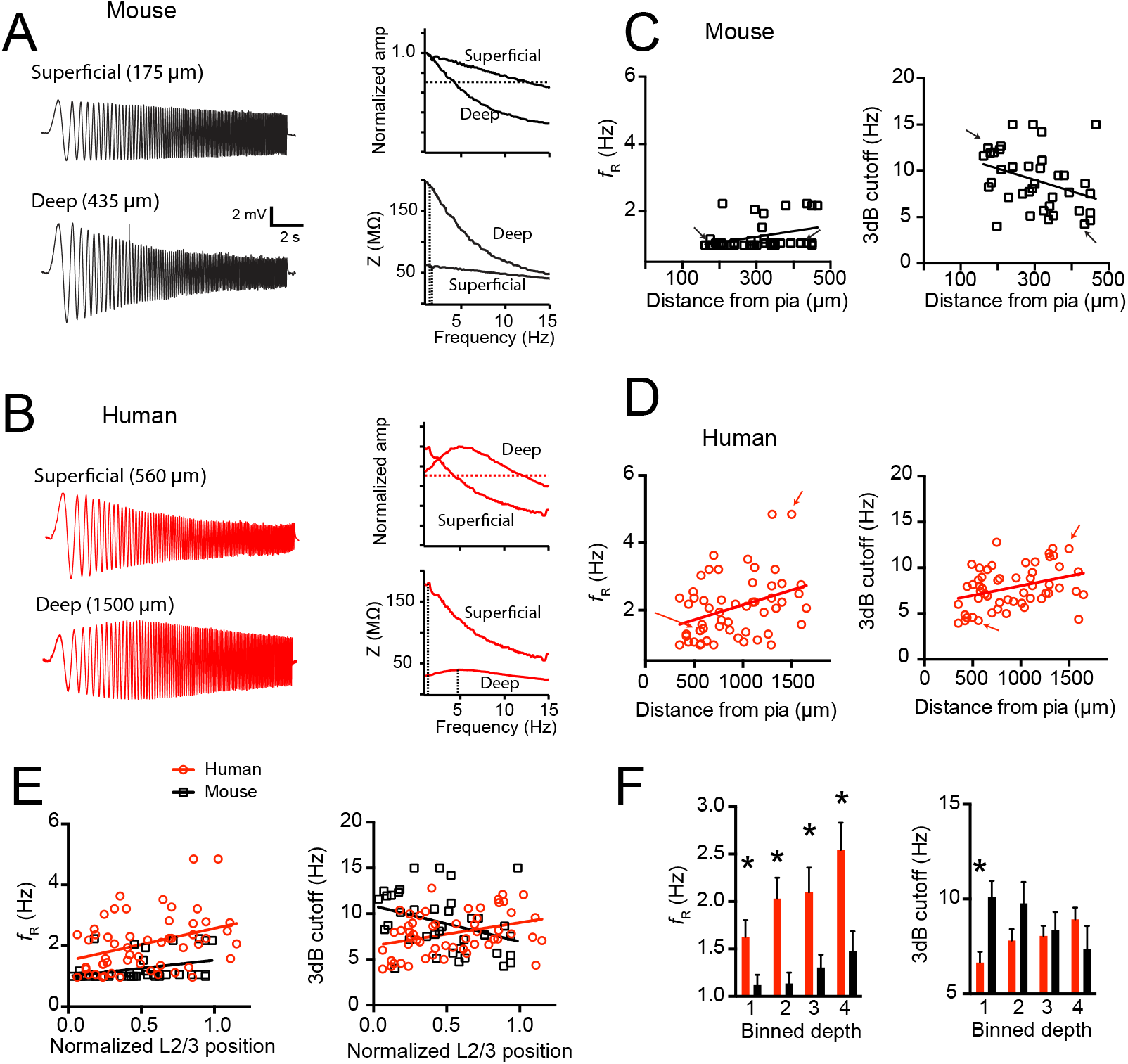
– Mouse and human supragranular pyramidal neurons display different subthreshold filtering properties. Example voltage responses to a chirp stimulus current injection in a superficial and deep supragranular pyramidal neuron in **A)** mouse and **B)** human cortex. ZAP and normalized frequency response curves are also shown for these example neurons. Dotted lines mark the resonant frequency in the ZAP and the 3dB cutoff in the normalized frequency response curves. **C)** Mouse neurons were largely non-resonant regardless of their position within supragranular cortex and became more low-pass as a function of somatic depth from pia. **D)** Resonant frequency correlated with somatic depth from pia in human cortex. Additionally, in human cortex, 3dB cutoff frequency was correlated with somatic depth from pia. Arrows correspond to sample voltage sweeps in A & B. **E)** Resonant frequency and 3dB cutoff as a function of normalized depth from pia in mouse and human supragranular cortex. **D)** Data binned into quadrants. For binned resonant frequency data * p < 0.001 mixed factor ANOVA effect of species. For binned 3 dB cutoff data * p < 0.0125, mouse versus human post-hoc t-test with Bonferroni correction.

The 3 dB cutoff and resonance frequency of mouse and human pyramidal neurons are also plotted as a function of normalized soma position in L2/3 (Figure 4E). Direct comparisons revealed that human neurons had a higher resonant frequency than mouse neurons at all relative distances from the pial surface (Figure 4F; p < 0.001; ANOVA). Furthermore, the most superficial human neurons had a lower 3dB cutoff than mouse neurons (Figure 4F; p = 0.01; ANOVA followed by post-hoc t test). These data highlight striking differences in the subthreshold filtering properties of mouse versus human L2/3 pyramidal neurons, further supporting the hypothesis that Ih is prominent in human, but not in mouse supragranular pyramidal neurons.

In addition to contributing to subthreshold properties, I_h_ impacts the action potential output of a neuron in response to suprathreshold current injections by affecting R_N_, RMP and after spike potentials (Brager and Johnston, 2007; Fan et al., 2005; Gu et al., 2005). Specifically, differences in the number of action potentials elicited by a given current injection can reflect differences in I_h_. Thus, we also tested for between-species differences in the response of individual pyramidal neurons to 1 s, depolarizing direct current injections (250, 500 and 750 pA). In mouse temporal association cortex, excitability (here defined as the number of spikes in response to a given current injection) was positively correlated with soma distance from the pial surface (Figure 5A; 250 pA r^2^ = 0.38, p < 0.001, 500 pA r^2^ = 0.17, p = 0.01, 750 pA r^2^ = 0.12, p < 0.03), such that the most excitable neurons were located deep in L3. In contrast to mouse cortex, excitability was negatively correlated with somatic distance from the pial surface in human cortex such that the most excitable neurons were located superficially (Figure 5B; 250 pA r^2^ = 0.21, p < 0.001, 500 pA r^2^ = 0.44, p < 0.001, 750 pA r^2^ = 0.47, p < 0.001). Example sweeps obtained from a superficial (top) and deep supragranular (bottom) mouse and human pyramidal neuron are shown in Figures 5A and B. Figure 5C plots the action potential frequency in response to three different amplitudes of current injection in mouse versus human temporal cortex normalized for the position of the recorded neuron through the depth of L2/3. Direct comparisons also revealed several depth-dependent differences in excitability between mouse and human pyramidal neurons (Figure 5D; ANOVA, p<0.001 followed by post-hoc t-tests). These general observations were replicated in a subset of experiments when the membrane potential was held at a common level via direct current injection (-65 mV, Figure S2 and 3). These data suggest that the depth-dependent intrinsic excitability of mouse and human pyramidal neurons differs dramatically. The most excitable human neurons were located superficially in supragranular cortex whereas the most excitable mouse neurons were located in the deepest part of the supragranular layers. While these results mirror the depth-dependent differences in Independent membrane properties described above, there are likely many contributors to these differences in excitability.

**Figure 5.**
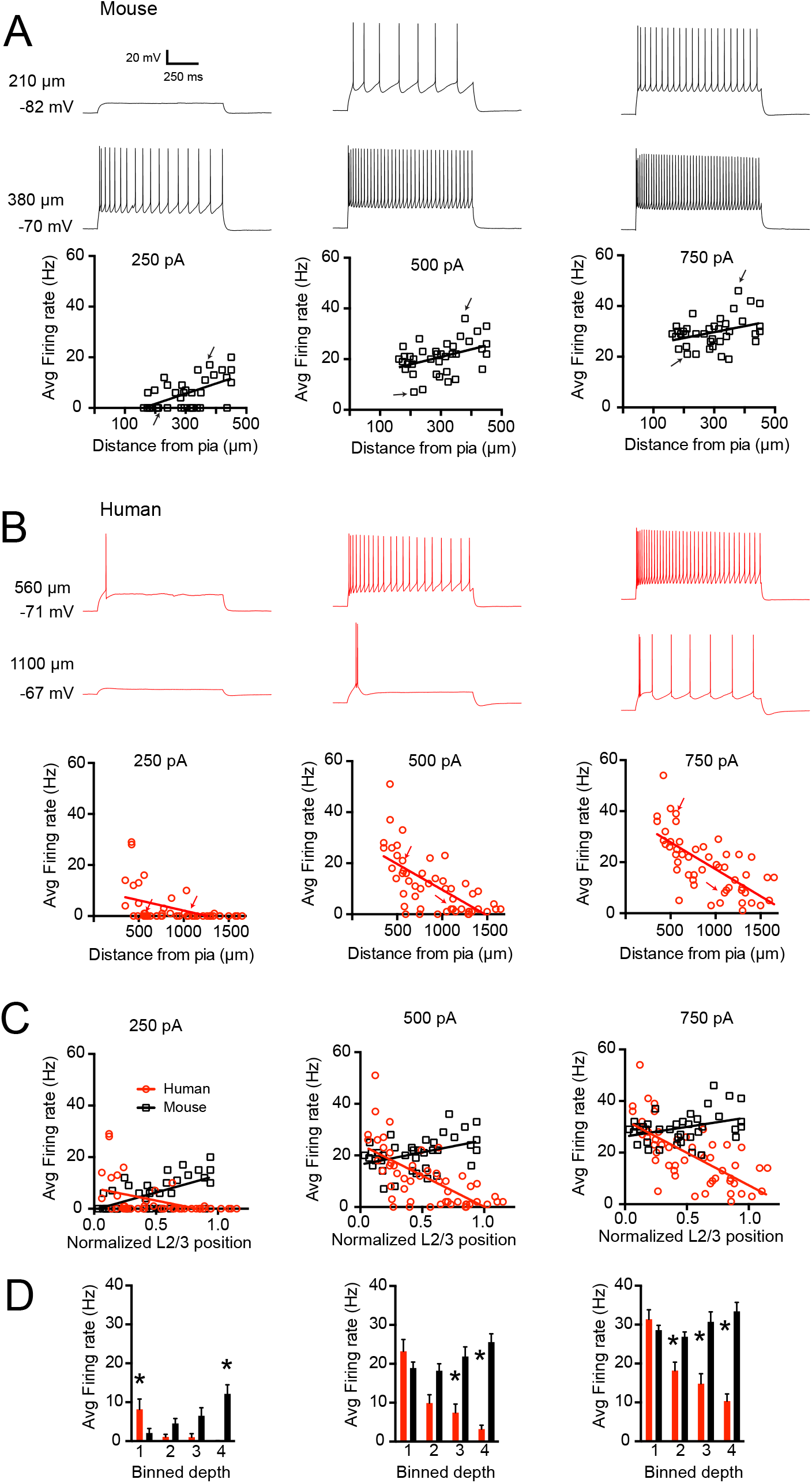
– Excitability of mouse versus human pyramidal neurons as a function of somatic distance from pia. **A)** The number of APs elicited by a given current injection increased as a function of somatic depth from pia in supragranular mouse cortex. Example sweeps obtained from a superficial and deep neuron in response to 250, 500 and 750 pA are shown. **B)** The number of APs elicited by a given current injection decreased as a function of somatic depth from pia in supragranular human cortex. Example sweeps obtained from a superficial and deep neuron in response to 250, 500 and 750 pA are shown. Arrows correspond to sample voltage sweeps in A & B. **C)** Average firing rate as a function of normalized position within supragranular cortex in mouse versus human. **D)** Average firing rate of mouse and human supragranular pyramidal neurons binned by somatic depth from pia. * < 0.0125 mouse versus human post-hoc t-test with Bonferroni correction.

The data presented thus far were collected from tissue obtained from patients with temporal lobe epilepsy. To assess the generality of the between-species differences in I_h_-related membrane properties, we also obtained data from temporal lobe tissue from a patient diagnosed with a deep brain tumor. The depth-dependent I_h_-related membrane properties observed in tissue obtained from epilepsy patients were also observed in supragranular cortex from this tumor patient (Figure S4). While we can’t rule out subtle differences, these data suggest that I_h_-related membrane properties in supragranular pyramidal neurons are not solely related to epilepsy.

### Pharmacological evidence for I_h_ in human supragranular pyramidal neurons

To test for the relative contribution of I_h_ to the intrinsic membrane properties of human versus mouse pyramidal neurons, we bath applied the h-channel blocker ZD7288 while monitoring resting membrane potential (Figure 6). For these experiments, we focused on recording from neurons located deeper in supragranular cortex, where I_h_-related properties were more apparent in both species. On average, ZD7288 produced a hyperpolarization of the RMP by 9.44 ± 0.80 mV in human neurons and -0.19 ± 1.68 mV in mouse neurons (Figure 6A). In addition, input resistance, when measured at a common potential of -65 mV increased by 62.14 ± 11.81% in human neurons (from 40.30 ± 2.37 MΩ to 64.35 ± 3.23 MΩ) and by 19.47 ± 7.89% percent in mouse neurons (from 130.63 ± 5.74 MΩ to 158.20 ± 11.86 MΩ; Figure 6B). ZD7288 also eliminated voltage sag, rebound and resonance, indicating that these membrane properties are dependent on functional h-channels in human supragranular pyramidal neurons (Figure S5). In addition, ZD7288 reduced the cutoff frequency of human neurons more so than mouse (Figure S5). Finally, ZD7288 increased the excitability of human pyramidal neurons, as is apparent in the parallel shift of the average input/output curve in Figure 6C. Together, these data suggest that I_h_ contributes to the intrinsic membrane properties of human supragranular pyramidal neurons significantly more so than mouse neurons.

**Figure 6.**
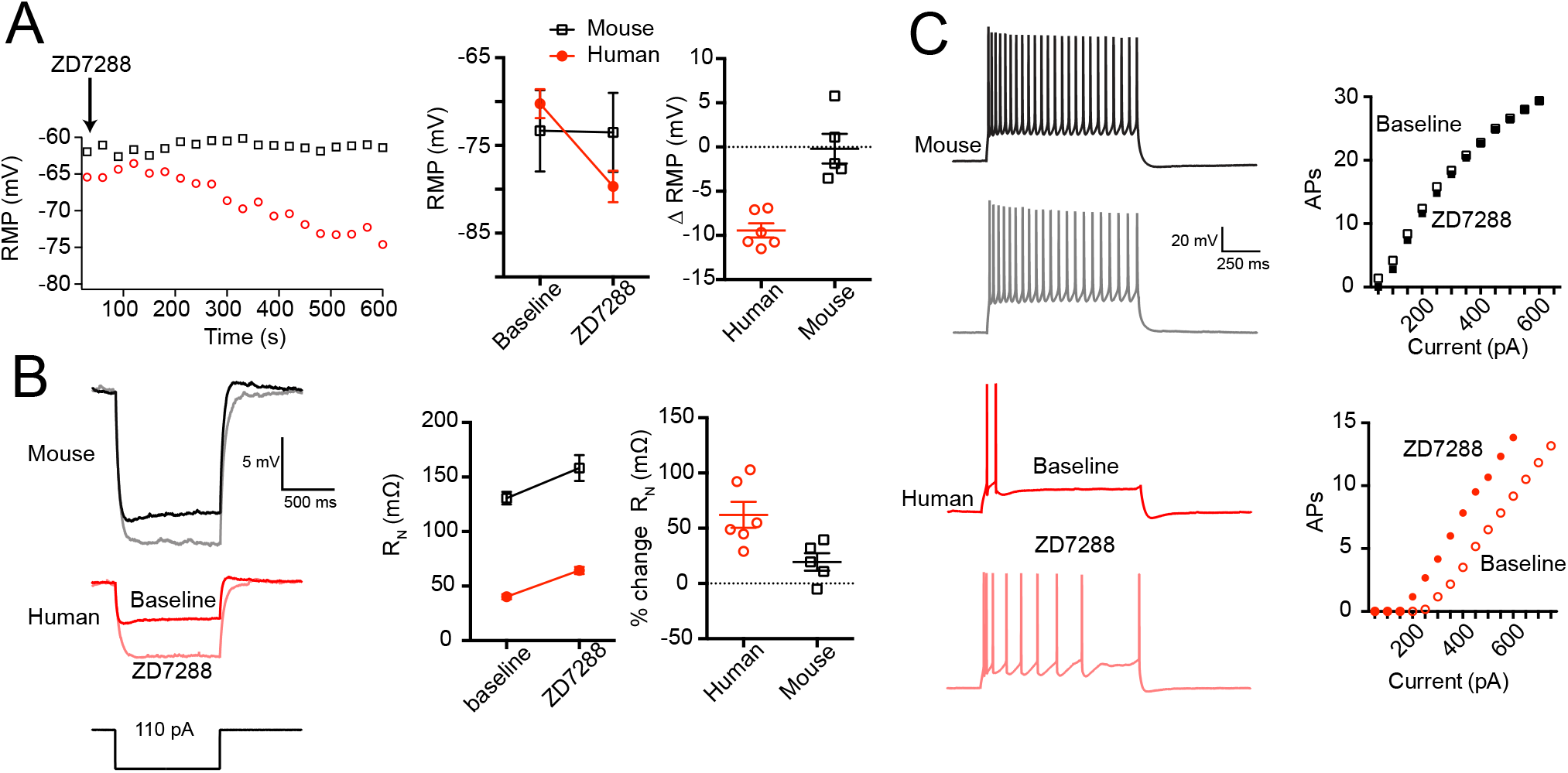
– Pharmacological evidence for I_h_ in human supragranular pyramidal neurons. **A)** Bath application of 10 μM ZD7288 produced a ~10 mV hyperpolarization of the resting membrane potential in human neurons, but no consistent change in mouse neurons. The plot at the left shows resting membrane potential as a function of time for two example recordings. **B)** 10 μM ZD7288 had a larger effect on the input resistance of human compared with mouse supragranular pyramidal neurons. Example voltage responses to hyperpolarizing current injections are shown to the left. **C)** Bath application of 10 μM ZD7288 increased the number of action potentials produced by a given current injection in human pyramidal neurons, but not mouse pyramidal neurons. Plots are averages from 6 human and 5 mouse experiments. Voltage responses are from example experiments.

### Impact of I_h_ on subthreshold integration in human neurons

What influence might I_h_ have on the input/output properties of human supragranular pyramidal neurons? While I_h_ affects many aspects of neuronal function, perhaps its most consistently observed influence is on subthreshold synaptic integration. h-channel expression in many neurons counteracts the distance-dependent capacitive filtering of synaptic input as it propagates from dendrite to soma (Dembrow et al., 2015; Harnett et al., 2015; Koch et al., 1990; Magee, 1999; Magee and Cook, 2000; Rall, 1967; Stuart and Spruston, 1998; Vaidya and Johnston, 2013; Williams and Stuart, 2000); this ensures that the kinetics of synaptic potentials at the soma are relatively independent of synaptic origin. Additionally, I_h_ narrows the window for temporal integration of synaptic input often to synaptic inputs with frequency components in the theta (4-12) band (Das and Narayanan, 2014; Dembrow et al., 2015; Kalmbach et al., 2017; Narayanan and Johnston, 2008; 2007; Ulrich, 2002; Vaidya and Johnston, 2013). To explore whether I_h_ might similarly affect the integrative properties of human supragranular pyramidal neurons, we used a morphologically precise (morphology is shown in Figure 7C) computational model of a human layer 3 pyramidal neuron (Methods) that possessed no active conductances other than I_h_. We chose to model a deep L3 pyramidal neuron where I_h_-related properties were more apparent.

**Figure 7.**
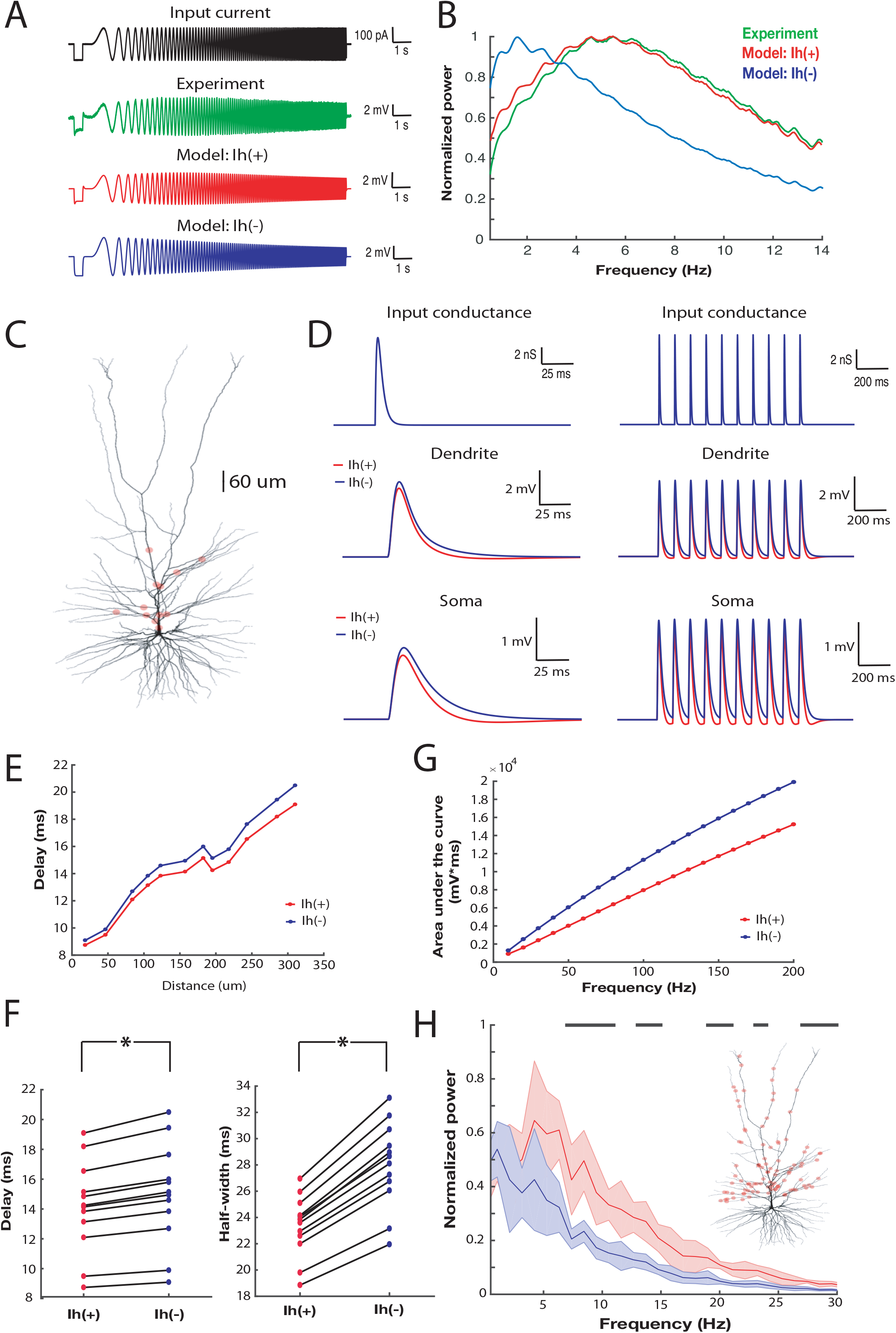
– –I_h_ affects the subthreshold integrative properties of a morphologically precise human L3 pyramidal neuron model. **A)** Intracellular somatic chirp stimulation produced a subthreshold somatic voltage response in a L3 pyramidal neuron (green). The identical stimulation protocol was imposed on a biophysically detailed model that includes (Ih(+); red) or excludes (I_h_(-); blue) I_h_. **B)** Power spectrum of somatic membrane potential response to intracellular chirp stimulation shown in panel A (blue: experiment; green: I_h_(+) model; red: I_h_(-) model). In both the I_h_(+) model and the experiment, the frequency response curves are nearly overlapping with a resonant peak at ~ 5Hz. **C)** Morphological reconstruction of a human L3 pyramidal neuron used for the simulations shown here **D)** Single (left) or bursts (right) of AMPA-like conductances were injected at single synaptic locations (top) and the resultant local dendritic and propagated somatic voltage response were recorded in the I_h_(+) (blue) and the I_h_(-) model (red). The locations of 12 separate synaptic inputs are shown in panel C). **E)** The delay between the maximal amplitude of AMPA-like conductance and EPSPs peak recorded at the soma as a function of synaptic distance from soma in the I_h_(+) (blue) and the I_h_(-) model. **F)** Synaptic delays and half-width of the EPSPs calculated for Ih(+) and Ih(-) models. G) The integral of EPSPs recorded at the soma in response to bursts of synaptic input at various frequencies. The somatic response in the I_h_(+) model was decreased relative to the I_h_(-) model across several frequencies of synaptic input. **H)** – Power spectrum of the somatic membrane potential of the I_h_(+) and I_h_(-) model when stimulated by 1000 synapses randomly located along the apical dendrite (synaptic conductance: 0.001 mS/cm^2^; morphology same as in panel C). Black stripes at 1 correspond to statistically significant differences in the power spectrum (2 sample Kolmogorov-Smirnov test; p<0.01). Inset: location of a subset (100 out of 1000) synapses is shown.

We first asked whether the presence of I_h_ in the model reproduced the subthreshold resonance observed in many deep supragranular pyramidal neurons in human temporal cortex (Figures 7A and B). The presence of I_h_ significantly affected the response of the model to a somatic chirp current injection. Notably, even though the chirp stimulus was not used to generate the computational model, in the presence of I_h_ (model: I_h_ (+)), the model displayed band pass filtering properties closely resembling those observed experimentally (Figures 7A and B). In contrast, in the absence of I_h_ (model: I_h_ (-)) the frequency response of the model markedly departed from the experimentally measured one (Figures 7A and B).

We next assessed how the presence of I_h_ in the model affected the integration of synaptic input arriving at various locations along the dendrite. To this end, we activated AMPA-like conductances at several locations along the dendritic arbor and measured the resultant local dendritic and propagated somatic voltage responses (Figures 7C and D). In a totally passive neuron, low-pass filtering severely attenuates and distorts synaptic inputs as they propagate to the soma, especially those arriving at distal locations (Koch, 2004; Koch et al., 1990; Rall, 1967; Stuart and Spruston, 1998). To quantify the effect of I_h_ on single EPSP kinetics, we measured the delay in the peak of the somatic EPSP relative to the peak of the local dendritic EPSP as well as the halfwidth of the somatic EPSP. In the passive model (I_h_(-)) a distance-dependent increase in the delay of the somatic EPSP relative to the dendritic synaptic conductance was observed as well as in the halfwidth of the somatic EPSP. In comparison, in the I_h_(+) model the delay between the peak of the dendritic and somatic EPSPs was significantly reduced, especially at distal locations, as was the halfwidth of the somatic EPSP (Figures 7E and F). Thus, the inclusion of I_h_ produced EPSPs at the soma of the model layer 3 human pyramidal neuron with a significantly faster time course.

These effects suggest that I_h_ influences the temporal summation of synaptic input in human supragranular pyramidal neurons; faster EPSP kinetics modestly reduce the temporal window wherein inputs can summate (Dembrow et al., 2015; Magee, 1999; Vaidya and Johnston, 2013; Williams and Stuart, 2000). To examine this possibility, we initiated bursts of AMPA-like conductances at various frequencies and different locations along the dendrite and measured the resulting somatic response. The total somatic depolarization (as quantified by the integral of the somatic voltage response) was reduced across several frequencies of synaptic input (Figure 7G). Thus, the presence of I_h_ in the model L3 human pyramidal neuron reduced the temporal summation of synaptic inputs.

Finally, by opposing changes to membrane potential, I_h_ can impart phenomenological inductance to the membrane. This has the effect of counteracting lags in the phase of membrane potential relative to current that is imposed by capacitive elements of the membrane (Koch, 1984; Mauro, 1961). This inductive property of I_h_ is also known to promote the transfer to the soma of synaptic input containing of theta frequencies (Cook et al., 2007; Dembrow et al., 2015; Narayanan and Johnston, 2008; Ulrich, 2002; Vaidya and Johnston, 2013). To test whether in human supragranular pyramidal neurons I_h_ also promotes the selective transfer of frequencies in the theta frequency range, we initiated AMPA-like synaptic input with a Poisson process (4 Hz) at 1000 locations along the dendritic arbor (Figure 7H-inset). Comparing the I_h_(+) and I_h_(-) models revealed that the presence of I_h_ resulted in an increase in power in the 5-15 Hz range of the somatic voltage response (Figure 7H). Thus, although the dendrite was presented with a random spatial-temporal pattern of input, frequencies in the 5-15 Hz range were preferably passed to the soma in the I_h_(+) model. The high pass filtering properties of I_h_ together with the low pass filtering properties of the passive dendritic membrane impart the band pass shape of this transfer function (Hutcheon and Yarom, 2000).

## Discussion

We have provided new evidence indicating a disparate contribution of h-channels to human versus mouse supragranular pyramidal neuron properties. In contrast to mouse, human supragranular excitatory neurons ubiquitously expressed *HCN1* transcripts as well as transcripts for *PEX5L*, a gene coding for an important regulatory protein of h-channel function. Consistent with this observation, we observed more pronounced Ih-related membrane properties in human compared with mouse pyramidal neurons at all distances from the pial surface, as well as depth-dependent differences in R_N_ and excitability. Furthermore, the h-channel blocker, ZD7288, affected these intrinsic membrane properties in human more so than in mouse supragranular pyramidal neurons. Finally, we used a computational model to provide evidence that the expression of h-channels in human supragranular pyramidal neurons narrows the window for synaptic integration by accelerating EPSP kinetics and promotes the transfer of synaptic input containing theta frequencies from the dendrite to soma.

Previous studies have largely focused on differences in the morphological, synaptic and/or passive membrane properties of rodent versus human supragranular pyramidal neurons. For example, human neurons are larger and possess a more complex dendritic arbor compared with mouse (Deitcher et al., 2017; Mohan et al., 2015). Differences in dendritic morphology and passive dendritic membrane properties could contribute to differences in the cable properties of human versus rodent pyramidal neurons (Eyal et al., 2016). Likewise, these differences, together with differences in synaptic properties may contribute to the reported high bandwidth processing capabilities of human neurons compared with mouse (Eyal et al., 2014; Testa-Silva et al., 2014). Our observation of cortical depth-dependent differences in mouse versus human pyramidal neuron properties adds to this growing list of interspecies disparities in cortical pyramidal neuron properties. Intriguingly, the depth-dependent differences in intrinsic properties we observe in human cortex parallel the lamination and differentiation of cell size that occurs along the radial axis of the supragranular layers (von Economo and Koskinas, 2007; Hill and Walsh, 2005; Molnár et al., 2014; Rakic, 2009). As such, the depth-dependence of these membrane properties may reflect an evolutionary adaptation for the expanded supragranular cortex of humans.

Our findings contrast with a previous report that found few correlations between L2/3 membrane properties and somatic depth from the pial surface in human cortex (Deitcher et al., 2017). This discrepancy may be due to differences in sampling. The dataset included in the previous report was smaller than the current set (n = 25 versus n = 55). Thus, it is possible that our data set captured more depth-dependent variability in intrinsic properties simply because of increased sampling. Relatedly, we sampled a wider range of somatic depths from the pial surface than the previous study (350-1600 μm in the current study; 471-1192 μm in the previous study). L2 begins ~275 μm from the pial surface and is ~ 150 μm wide in human temporal cortex (von Economo and Koskinas, 2007). Similarly, deep L3 can extend up to 1600 μm from the pial surface in temporal cortex (von Economo and Koskinas, 2007). Thus, the previous study may not have sampled the most superficial of L2 or the deepest portions of L3, where the largest differences in intrinsic membrane properties exist.

To our knowledge our findings are the first to directly implicate a particular ion channel in differences between human and rodent pyramidal neuron properties. Supragranular pyramidal neurons in rodent cortex express little h-channel-related protein or RNA (Figure 1; Lörincz et al., 2002; Santoro et al., 2000; Zeng et al., 2012). Furthermore, mouse and rat supragranular pyramidal neurons display very few hallmarks of I_h_ (e.g. sag) across several areas of cortex (Larkum et al., 2007; Routh et al., 2017; van Aerde and Feldmeyer, 2013). Together, these observations suggest that I_h_ contributes very little to rodent supragranular pyramidal neuron physiology regardless of cortical region. In contrast, human supragranular pyramidal neurons display prominent voltage sag (Deitcher et al., 2017; Foehring and Waters, 1991) that we show here is dependent upon h-channels. Furthermore, sag and other I_h_-related properties in human cortex are more prominent in deep, compared to superficial, L2/3 pyramidal neurons. We note, however, that I_h_ is unlikely to explain all of the depth-dependent differences in intrinsic properties we observed between mouse and human pyramidal neurons. Clearly other factors, including in morphology and/or differential expression of conductances other than I_h_ may contribute to the depth- and species-dependent differences in excitability, R_N_ and subthreshold filtering we observed.

Our findings have a few limitations and caveats that are inherent to studying human brain tissue at this level of analysis. First, human physiology data were collected from tissue obtained from neurosurgical patients and thus may be influenced by neurological disease state. Notably, h-channels are implicated in epilepsy (Brennan et al., 2016; Jung et al., 2007; Shin et al., 2008) and thus our results may be influenced by neuropathology. Several factors, however, strengthen our conclusion that Ih prominently contributes to the membrane properties of human supragranular pyramidal neurons in the neurotypical condition. First, the tissue obtained for these experiments was distal to the focus of the seizures and did not express overt signs of pathology. Second, we found similarly prominent I_h_-related properties in 23 supragranular pyramidal neurons of temporal cortex brain slices (Figure S4) derived from tumor patients and in other cortical regions (Allen Cell Types data base - http://celltypes.brain-map.org; Deitcher et al., 2017). Finally, h-channel subunit RNA is abundant in pyramidal neurons in the supragranular layers of cortical post-mortem tissue obtained from donor brains with no prior history of neurological disorder (Figure 1; Zeng et al., 2012).

### Functional implications

Deep L3 neurons possess the most prominent I_h_-related membrane properties in the supragranular layers of human temporal cortex. Deep L3 also corresponds to the sub-lamina containing the largest supragranular pyramidal neurons with the largest total dendritic length (von Economo and Koskinas, 2007; Mohan et al., 2015). Because of their large size, these neurons may require specialized intrinsic mechanisms to ensure faithful propagation of signals along their dendritic arbor. Our modeling results suggest that h-channels may serve in this regard by counteracting capacitive filtering by the dendrite.

Our simulations also demonstrate that h-channels significantly affect the integrative properties of human supragranular pyramidal neurons in a similar manner as has been reported in rodent cortical L5 or hippocampal CA1 pyramidal neurons. (Dembrow et al., 2015; Vaidya and Johnston, 2013; Williams and Stuart, 2000). Consistent with its role as a resonating conductance (Hu et al., 2002; Hutcheon and Yarom, 2000; Narayanan and Johnston, 2008), the presence of I_h_ resulted in an increase in power in the theta (and higher) frequency range (Figure 7H). The specific details of these effects will depend upon several factors, including total abundance, subcellular localization and/or gradients of the channels. In rodents, h-channels are enriched in the distal apical dendrites of several types of pyramidal neurons in multiple brain regions (Harnett et al., 2015; Kalmbach et al., 2013; Kole et al., 2006; Lörincz et al., 2002; Williams and Stuart, 2000). Notably, we observed widespread expression in human temporal cortex of a gene (PEX5L) that codes for Trip8b, a protein that is necessary for dendritic enrichment of I_h_ (Lewis et al., 2009; 2011). Thus, our results are consistent with the possibility that I_h_ is prominent in the dendrites of human supragranular pyramidal neurons. Nevertheless, modeling studies suggest that the effects of I_h_ on EPSPs kinetics do not appear to depend on subcellular gradients, but rather on total expression levels (Angelo et al., 2007; Das and Narayanan, 2014). Thus, the effects of I_h_ we observed on synaptic integration in our single neuron model may not depend upon the exact localization of h-channels, but rather, their total expression levels.

Finally, our results suggest that I_h_ may significantly affect the spike initiation dynamics of human supragranular pyramidal neurons. The presence of h-channels can switch the firing mode of a neuron from temporal integrator to coincidence detector, whereby spiking is sensitive to correlated synaptic input rather than changes in mean presynaptic firing rate (Das and Narayanan, 2017; 2014; Ratté et al., 2013). For rodent neurons, there is an intimate relationship between sub- and suprathreshold spectral selectivity (Das and Narayanan, 2017; 2014). Thus, the spiking activity of human supragranular pyramidal neurons may be tuned to specific frequencies of synaptic input, and this selectivity may vary with somatic depth from the pial surface. If so, the expression of h-channels in supragranular pyramidal neurons may contribute to memory-related theta-frequency phase-locking of single human neurons observed *in vivo* (Jacobs et al., 2007; Rutishauser et al., 2010).

## Acknowledgments

We wish to thank the Allen Institute founder, Paul G. Allen, for his vision, encouragement and support. We also wish to thank the Allen Institute for Brain Science Tissue Procurement, Tissue Processing and Facilities teams for help in coordinating the logistics of human tissue surgical tissue collection, transport and processing. We also thank the Allen Institute for Brain Science Reagent Preparation, Animal Care and In Vitro Single Cell Characterization teams. We are grateful to our collaborators at the local hospital sites, including Tracie Granger, Caryl Tongco, Matt Ormond, Jae-Guen Yoon, Nathan Hansen, Niki Ellington, Rachel Iverson (Swedish Medical Center), Carolyn Bea, Gina DeNoble and Allison Beller (Harborview Medical Center). We thank Dirk C. Keene at Harborview/UW for consultation and support. We also thank Meanhwan Kim for helpful discussions and suggestions.

